# INHIBITION OF LIGAND-DEPENDENT BMP SIGNALING BLUNTS MELANOMA GROWTH

**DOI:** 10.64898/2026.06.25.734518

**Authors:** Alec K. Gramann, Monir Ejemel, Arvind M. Venkatesan, Lindsay M. Ferreira, Carey Zammitti, Danielle Wiseheart, Yang Wang, Michael Brehm, Craig J. Ceol

## Abstract

Treatments for advanced melanoma have markedly improved, but a significant proportion of patients still receive little to no survival benefit with standard-of-care therapies due to resistance and relapse^1–5^. The discovery and development of novel targets and therapies are needed to continue to improve patient outcomes in advanced melanoma. The identification of ligand-dependent BMP signaling that inhibits differentiation and promotes survival of melanoma cells suggests it is a potential therapeutic target that could complement current therapies^6^. Expression of the BMP ligand GDF6 (a.k.a BMP13) is responsible for this activity, and its expression is correlated with poor outcomes for melanoma patients. Here, we describe a novel monoclonal antibody targeting GDF6 that causes melanoma cell differentiation and death and blunts tumor growth in vivo. Together, these results indicate BMP-directed therapy has significant potential as a novel therapy for patients with advanced melanoma.

## RESULTS & DISCUSSION

Previous studies found that BMP signaling is evident in over 70% of melanomas, and it is activated by the tumor-expressed BMP ligand GDF6^6,7^. To target this activity in a way that would limit damaging on-target effects of broadly inhibiting BMP signaling in normal tissues, we aimed to generate antibodies that would target GDF6, loss of which affects embryonic bone, cochlear and melanocyte development but does not impact adult tissue growth or function^8–12^. Whereas most BMP ligands are secreted and act in the extracellular space, it has been shown in specific contexts that BMP ligands can have limited diffusion capacity and be sequestered near the surface of cells, potentially limiting their targetability in extracellular space^13–15^. We first confirmed that GDF6 was indeed secreted and able to be isolated away from the surface of melanoma cells by collecting conditioned media and probing for the presence of GDF6 using an anti-GDF6 antibody (Figure 1A). To confirm that this extracellular GDF6 was able to bind the surface of melanoma cells where BMP receptors are expressed, we developed a cell-surface binding assay using hemagglutinin-tagged GDF6 (GDF6-HA) (Figure 1B). GDF6-HA was isolated from conditioned media and added to media of parental A375 melanoma cells. We analyzed the cells for surface binding of GDF6-HA by flow cytometry. Cells treated with GDF6-HA-enriched media, or cells expressing GDF6-HA endogenously, showed cell-surface binding of the tagged protein (Figure 1C). We wanted to verify that exogenous GDF6 provided by conditioned media was able to functionally act on melanoma cells. We previously showed knockdown of GDF6 caused significant loss of viability in A375 and other melanoma cell lines^6^. We knocked down GDF6 in A375 cells and treated with conditioned media with depleted, normal, and enriched levels of GDF6 protein and assessed cell viability. The treatment of GDF6-knockdown cells with media containing normal or enriched levels of GDF6 protein rescued the viability, whereas treatment with GDF6-depleted media failed to rescue (Figure 1D). Together, these results indicate GDF6 is secreted by melanoma cells and is capable of promoting melanoma cell viability when provided exogenously.

**Figure 1.**
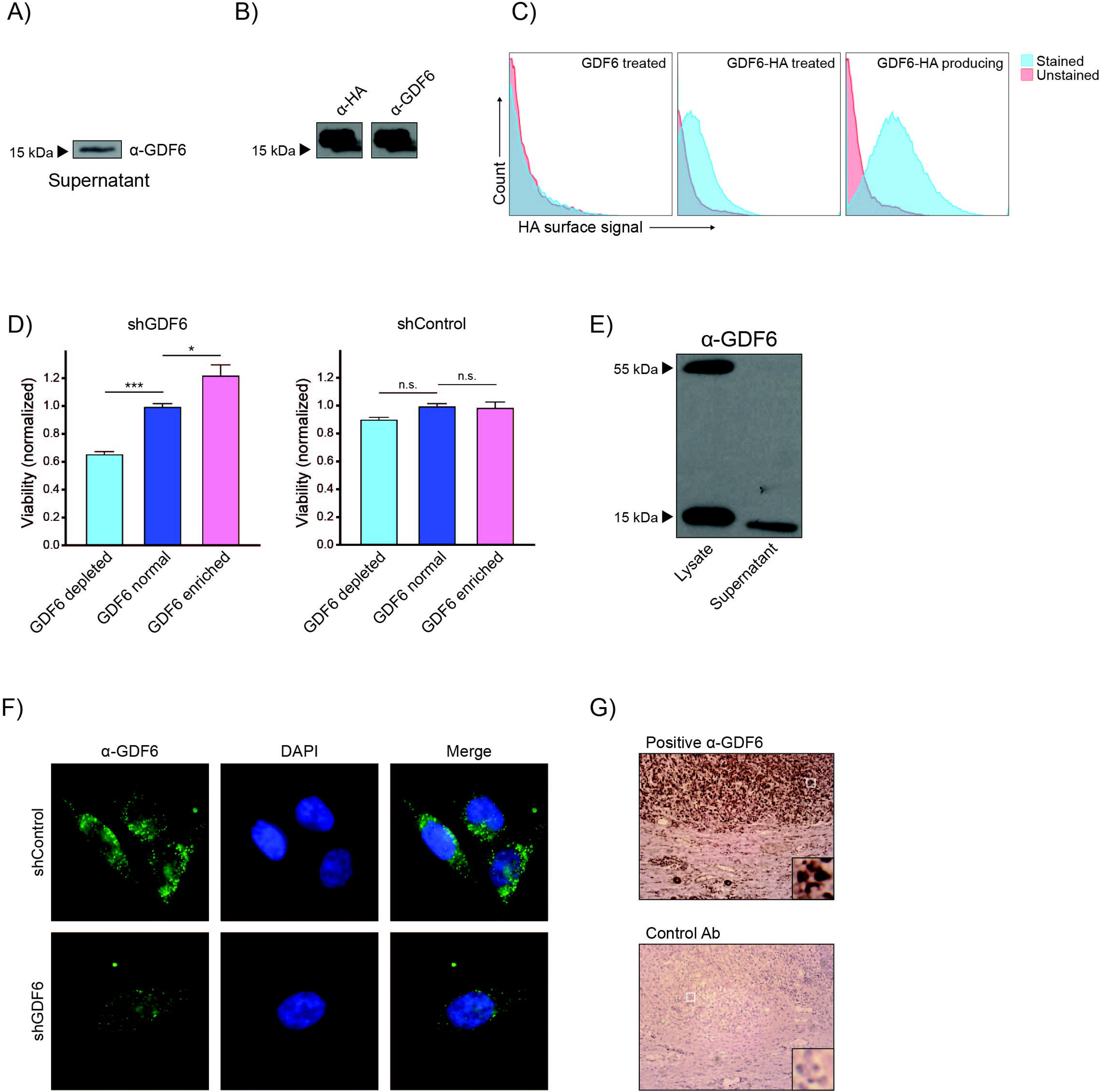
GDF6 is secreted and functions extracellularly on melanoma cells. (A) Western blot of conditioned media from A375 melanoma cells showing the 15 kDa mature form of GDF6. (B) Western blot of HA-tagged GDF6 from conditioned media and identified by anti-HA antibody and anti-GDF6 antibody. (C) Flow cytometry histograms showing increased GDF6-HA surface signal on A375 cells treated with conditioned media containing GDF6-HA or on cells producing GDF6-HA. Unstained control samples were not stained with Alexafluor-488-conjugated anti-HA antibody. (D) Relative viability of A375 cells expressing shControl (shEGFP) or shGDF6 when treated with GDF6-depleted, GDF6-normal, and GDF6-enriched conditioned media. Cells were treated with GDF6 media for 24 hours prior to assessing viability. n = 4-6 technical replicates across two independent experiments (N=2). (E) Anti-GDF6 mAb specificity for GDF6 protein isolated from A375 melanoma cell lysates and conditioned media shows recognition of full-length 55 kDa pro-GDF6 and 15 kDa mature GDF6. (F) Immunofluorescence using anti-GDF6 mAb showed a decrease in fluorescence intensity in cells expressing shGDF6 relative to shControl. (G) Immunohistochemical staining of patient tumor samples using anti-GDF6 mAb and a control antibody that did not recognize GDF6 (brown pigment) in bulk tumor. Error bars represent mean +/- SEM. P-values were calculated for panel D using one-way ANOVA with Tukey’s multiple comparisons correction, * p<0.05, *** p<0.001, n.s., not significant.

As a factor that acts extracellularly, GDF6 is amenable to targeting by a monoclonal antibody to bind and block the ability of GDF6 to activate BMP signaling. To develop anti-GDF6 monoclonal antibodies, we immunized mice with MBP-GDF6C, a recombinant protein fusion of maltose binding protein and mature GDF6 protein, which is made up of the C-terminal 120 amino acid of the GDF6 proprotein. Hybridoma clones from these mice were screened by ELISA for activity against the GDF6 immunogen, and ELISA-positive clones were selected for further evaluation (Supplementary Figure 1A). ELISA-positive antibodies were screened by western blot for their specific recognition of native GDF6 (Supplementary Figure 1B), and the effects of antibodies on melanoma cell viability were tested (Supplementary Figure 1C). We describe one of these anti-GDF6 monoclonal antibodies, hereafter referred to as anti-GDF6 mAb, identified from this initial screen. Following purification of anti-GDF6 mAb from hybridoma supernatant, we assessed its ability to recognize GDF6 produced by melanoma cells. We found in western blots that the anti-GDF6 mAb exhibited high specificity for GDF6, binding both the full-length 55 kDa GDF6 proprotein and the 15 kDa mature, secreted form of GDF6 from A375 cell lysate (Figure 1E). In conditioned media from A375 cells, the anti-GDF6 mAb recognized the mature, secreted form only. We performed immunofluorescence using anti-GDF6 mAb in GDF6 knockdown cells and observed a decrease in fluorescence intensity, further indicating specificity of our anti-GDF6 mAb for GDF6 (Figure 1F). Immunohistochemical staining of patient melanomas showed staining by anti-GDF6 mAb in the bulk tumor (Figure 1G), consistent with previous observations that GDF6 expression is tumor-specific^6^. Together, these results show our selected anti-GDF6 mAb is able to bind human GDF6 in melanoma cells, both in vitro and in patient samples.

Knockdown of GDF6 in melanoma cells was previously shown to decrease canonical BMP signaling, leading to upregulation of a melanocytic differentiation program and pro-apoptotic factors^6^. Thus, we hypothesized treatment with anti-GDF6 mAb should result in similar pathway and transcriptional changes. As a positive control for changes to BMP signaling in melanoma cells, we used the small molecule inhibitor of BMP signaling, DMH1, hereafter referred to as BMPi^16^. We performed immunofluorescence for the marker of BMP activity, phospho-SMAD-1/5/8, in cells treated with anti-GDF6 mAb. We observed lower nuclear fluorescence intensity in cells treated with anti-GDF6 mAb (Figure 2A), which was comparable to the level in cells treated with BMPi, indicating anti-GDF6 mAb treatment was effective at blocking BMP signaling in melanoma cells. We found a similar decrease in phospho-SMAD-1/5/8 compared to control by western blot (Supplementary Figure 2A,B). To determine if this decrease in phospho-SMAD-1/5/8 intensity correlated with decreased transcriptional regulatory activity, we employed a luciferase construct driven by tandem BMP-binding elements isolated from the ID1 promoter, a canonical BMP target^17^. We observed similar decreases in luciferase activity in anti-GDF6 mAb- and BMPi-treated cells when compared to their respective controls (Figure 2B), further indicating anti-GDF6 mAb is capable of inhibiting GDF6-activated BMP signaling in melanoma cells. Some important targets in melanoma cells, *MITF* and *SOX9*, are normally repressed by GDF6-driven BMP signaling to inhibit differentiation and cell death, respectively^6,7,9^. RT-qPCR for these genes showed increased expression of both in response to treatment with anti-GDF6 mAb, as well as treatment with BMPi (Figure 2C,D). Together, these results indicate treatment with anti-GDF6 mAb produces similar changes to BMP signaling and transcriptional regulation as does GDF6 knockdown or treatment with a small-molecule BMP inhibitor.

**Figure 2.**
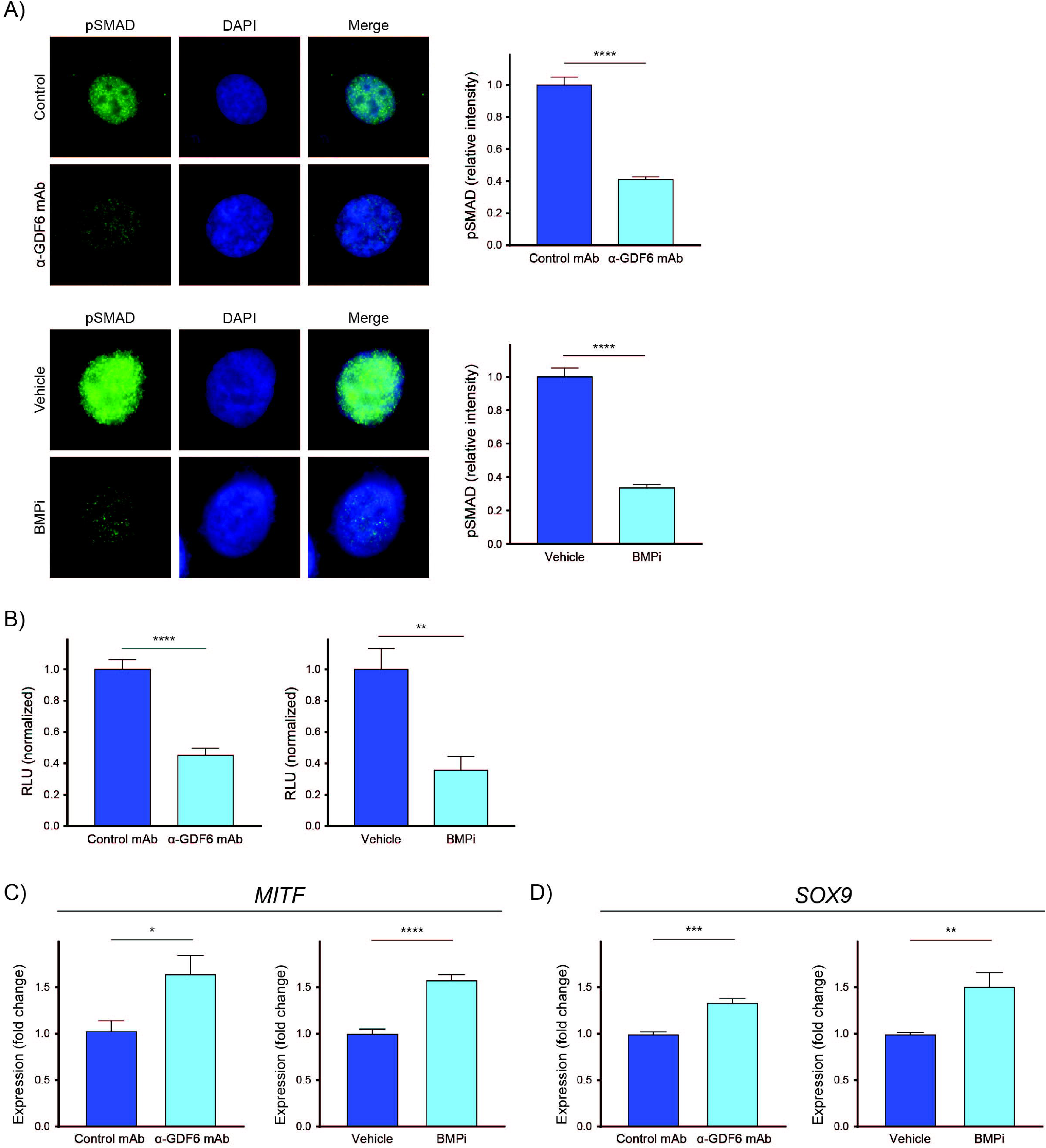
anti-GDF6 mAb blocks BMP signaling in vitro. (A) Top left, immunofluorescence for phospho-SMAD-1/5/8 (pSMAD) in SK-MEL-28 melanoma cells treated with control mAb or anti-GDF6 mAb. Top right, quantification of relative signal intensity of nuclear phospho-SMAD between control mAb and anti-GDF6 mAb-treated cells. n = 54 and 48 cells for control mAb and anti-GDF6 mAb groups, respectively, from two independent experiments (N = 2). Bottom left, immunofluorescence of pSMAD in SK-MEL-28 melanoma cells treated with vehicle or BMPi. Bottom right, quantification of relative signal intensity of nuclear pSMAD between vehicle- and BMPi-treated cells. n = 47 and 60 cells from vehicle and BMPi groups, respectively, from two independent experiments (N = 2). (B) Left, normalized relative luminescence units (RLU) in cells expressing BMP-responsive luciferase when treated with control mAb or anti-GDF6 mAb. n = 4-6 technical replicates across two independent experiments (N = 2). Right, RLU in cells expressing BMP-responsive luciferase when treated with vehicle or BMPi. n = 4-6 technical replicates across two independent experiments (N = 2). (C) qRT-PCR for *MITF* showing increase in expression in samples treated with anti-GDF6 mAb compared to control (left) and samples treated with BMPi compared to vehicle (right). n = 4-6 technical replicates across two independent experiments (N = 2). (D) qRT-PCR for *SOX9* showing increase in expression in samples treated with anti-GDF6 mAb compared to control (left) and samples treated with BMPi compared to vehicle (right). n = 4-6 technical replicates across two independent experiments (N = 2). Error bars represent mean +/- SEM. P-values calculated use Student’s t-test for panels B, D, E, F, G, H, * p<0.05, ** p<0.01, *** p<0.001, **** p<0.0001.

In our previous study, we found knockdown of GDF6 in cell lines with GDF6 amplification caused a BMP-dependent decrease in cell viability^6^. However, cell lines without GDF6 amplification were insensitive to GDF6 knockdown. We treated GDF6 knockdown-sensitive (A375, SK-MEL-28) and GDF6 knockdown-insensitive (M14, C32) melanoma cell lines with anti-GDF6 mAb and determined viability of the cells. We observed a decrease in viability of GDF6-sensitive cell lines and little change in viability of GDF6-insensitive cell lines upon treatment with anti-GDF6 mAb, with similar findings upon treatment with BMPi (Figure 3A). To determine if this GDF6-dependent decrease in viability was dependent on inhibition of BMP signaling, we performed anti-GDF6 mAb treatment on A375 cells expressing SMAD1-DVD, a constitutively active SMAD1 construct^18^. In cells expressing SMAD1-DVD, we observed a smaller loss of viability when compared to empty vector-expressing cells (Figure 3B), indicating the loss of viability was in part dependent on BMP signaling. We evaluated the longitudinal effect of anti-GDF6 mAb treatment on A375 melanoma cells over the course of 4 days of treatment. We observed a decrease in viability of cells treated with anti-GDF6 mAb over the course of treatment, compared to control mAb-treated cells, which exhibited an increase in viability (Figure 3C). In cells treated with BMPi, the cells exhibited lower viability than vehicle-treated control cells but had greater viability on day 4 as compared to the same cells on day 0, immediately prior to treatment. Comparison of the anti-GDF6 mAb and BMPi results suggested that a combination of mechanisms may be underlying the changes in viability by anti-GDF6 mAb treatment: a cidal effect -- decreasing viability by promoting cell death -- and a static effect -- decreasing viability by suppressing proliferation. To assess these effects we performed a caspase 3/7 activity assay and immunofluorescence staining for cleaved caspase-3 and Ki-67. A375 cells treated with anti-GDF6 had markedly increased caspase 3/7 activity (Figure 3D). Melanoma cells treated with anti-GDF6 mAb and BMPi showed both an increase in cleaved caspase-3 (Figure 3E) and a decrease in Ki-67 (Figure 3F), indicating both increased cell death and decreased proliferation contribute to the overall change in viability.

**Figure 3.**
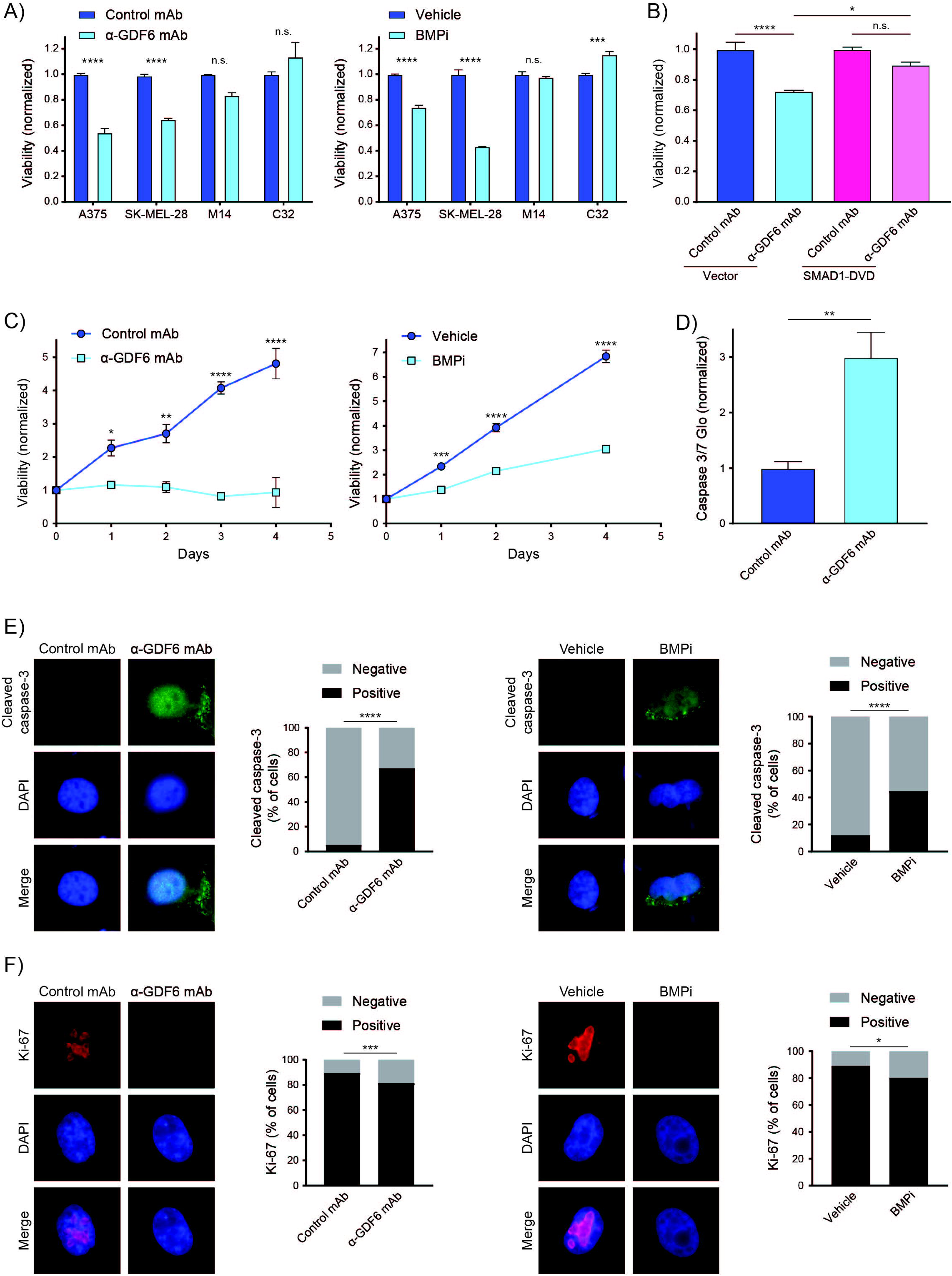
anti-GDF6 mAb decreases cell viability in a GDF6- and BMP-dependent manner. (A) Left, assessment of viability using CellTiter Glo of A375, SK-MEL-28, M14, and C32 melanoma cells following 24 hours of treatment with control mAb or anti-GDF6 mAb. n = 4-6 technical replicates across two independent experiments (N = 2). Right, assessment of viability of A375, SK-MEL-28, M14, and C32 melanoma cells following 24 hours of treatment with vehicle or BMPi. n = 4-6 technical replicates across two independent experiments (N = 2). (B) Assessment of viability of A375 cells expressing vector or SMAD1-DVD following treatment with control mAb or anti-GDF6 mAb. n = 4-6 technical replicates across two independent experiments (N = 2). (C) Left, longitudinal treatment of SK-MEL-28 melanoma cells with control mAb or anti-GDF6 mAb. Cells were treated on day 0 and viability was assessed daily by CellTiter Glo for 4 days. n = 4-6 technical replicates across two independent experiments (N = 2). Right, longitudinal treatment of SK-MEL-28 melanoma cells with vehicle or BMPi. Cells were treated on day 0 and viability was assessed daily by CellTiter Glo for 4 days. n = 4-6 technical replicates across two independent experiments (N = 2). (D) Normalized Caspase 3/7 Glo luminescence in cells treated with control mAb or anti-GDF6 mAb. n = 4-6 technical replicates across two independent experiments (N = 2). (E) Left, immunofluorescence and quantification of cleaved caspase-3 in SK-MEL-28 melanoma cells treated with control mAb or anti-GDF6 mAb. n = 923 and 621 cells for control mAb and anti-GDF6 mAb groups, respectively, from three independent experiments (N = 3). Right, immunofluorescence and quantification of cleaved caspase-3 in SK-MEL-28 melanoma cells treated with vehicle or BMPi. n = 291 and 145 cells for vehicle and BMPi groups, respectively, from two independent experiments (N = 2). (F) Left, immunofluorescence and quantification of Ki-67 in SK-MEL-28 melanoma cells treated with control mAb or anti-GDF6 mAb. n = 413 and 474 cells for control mAb and anti-GDF6 mAb groups, respectively, from two independent experiments (N = 2). Right, immunofluorescence and quantification of Ki-67 in SK-MEL-28 melanoma cells treated with vehicle or BMPi. n = 176 and 173 cells from vehicle and BMPi groups, respectively, from two independent experiments (N = 2). Error bars represent mean +/- SEM. P-values were calculated using one-way ANOVA with Tukey’s multiple comparison correction in panels A and B, Student’s t-test in panels C and D, and Fisher’s exact test in panels E and F, * p<0.05, ** p<0.01, *** p<0.001, **** p<0.0001, n.s., not significant.

Given the results showing anti-GDF6 mAb decreased melanoma cell viability in vitro, we wanted to assess the effectiveness of anti-GDF6 mAb treatment in an *in vivo* setting. Treatment with anti-GDF6 mAb of SK-MEL28 melanoma cell line xenografts in athymic nude mice resulted in an overall reduction in tumor volume and, in some animals, tumor regression was observed (Figure 4A, Supplementary Figure 3). Immunohistochemistry of tumor sections showed that anti-GDF6 mAb-treated tumors had increased cleaved caspase-3 positivity and decreased Ki67 (Figure 4B,C), mirroring results of treated cell lines in vitro. Additionally, immunohistochemistry found a reduction of phospho-SMAD1/5/8 in anti-GDF6 mAb-treated tumors, indicating an inhibition of BMP signaling in these tumors (Figure 4D). Together, these results indicate anti-GDF6 mAb therapy can inhibit GDF6-driven BMP signaling and blunt tumor growth in vivo.

**Figure 4.**
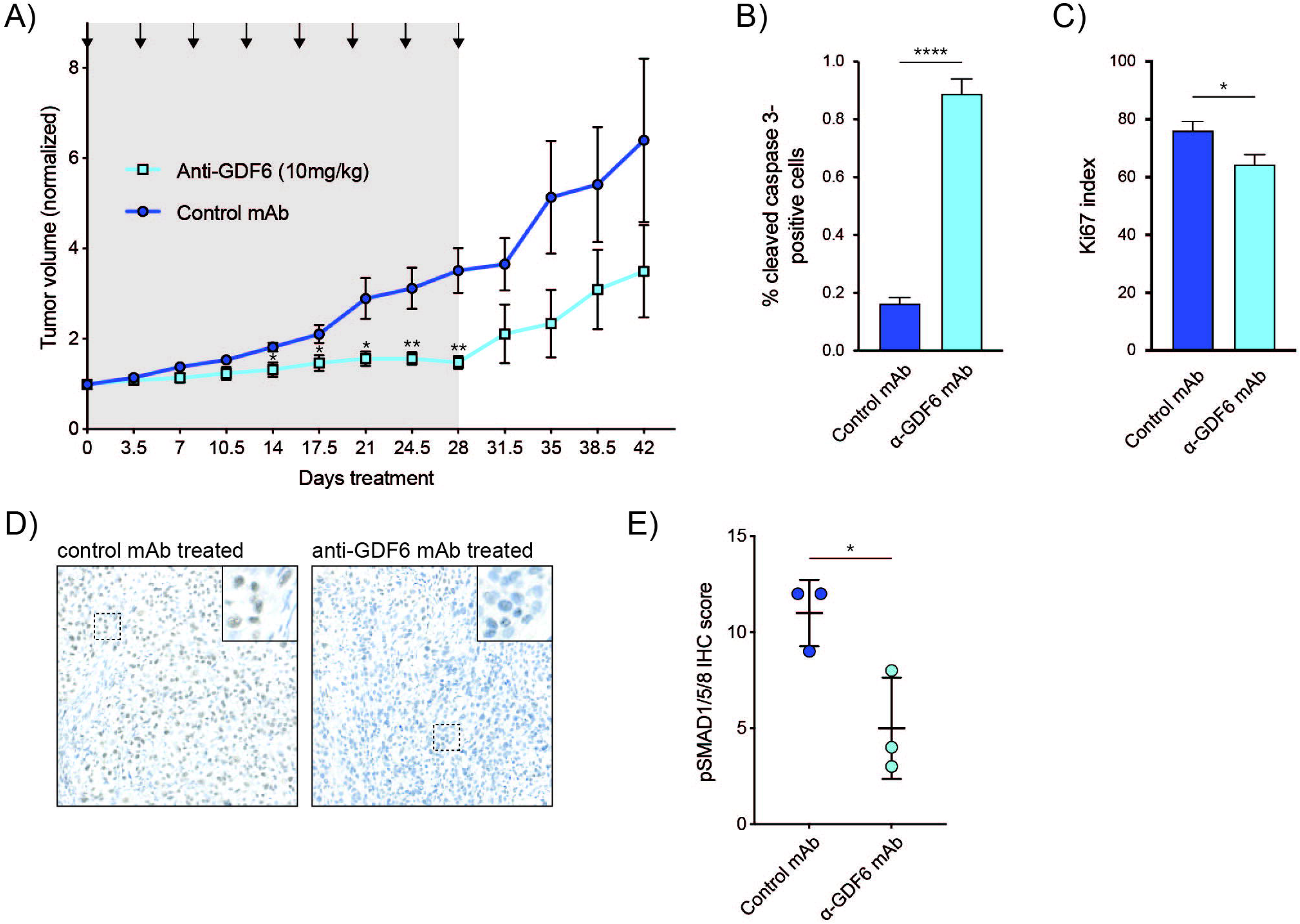
anti-GDF6 mAb blunts tumor growth in vivo. (A) Treatment of melanoma xenografted mice with anti-GDF6 mAb. Mice were injected with SK-MEL-28 cells and melanomas allowed to reach 100mm3, whereupon treatment with anti-GDF6 mAb was begun. n = 6 and 7 mice from control mAb and anti-GDF6 mAb groups, respectively. (B) Quantification of cleaved caspase 3 staining of xenografted tumor sections. n = 150 fields. (C) Quantification of Ki67 staining of xenograted tumor sections. n = 20 fields. (D) phosphorylated-SMAD1/5/8 immunohistochemistry of xenografted tumor sections. (E) Quantification of phosphorylated-SMAD1/5/8 using immunointensity x immunopositivity scoring of tumor nuclei. Error bars represent mean +/-SEM in panels A, B, C and +/- SD in panel E. P-values calculated use Student’s t-test for panels A, B, C, E, * p<0.05, ** p<0.01, **** p<0.0001.

Our study identifies a newly developed anti-GDF6 mAb that is a candidate for melanoma therapy. Anti-GDF6 mAb is able to bind GDF6 produced by human melanoma cell lines as well as human tumor samples. Treatment of melanoma cell lines with anti-GDF6 mAb is able to decrease BMP signaling activity, and leads to a corresponding decrease cell viability in a BMP-dependent manner. The change in viability by treatment with anti-GDF6 mAb is due to both an increase in cell death and a decrease in proliferation of melanoma cells. We observe similar findings in an in vivo setting, where treatment with anti-GDF6 mAb is able to drive regression of melanoma xenografts. These findings indicate anti-GDF6 mAb could be used to treat the 70-80% of human patient melanomas that express high levels of GDF6 to promote tumor regression. The mechanism by which it acts, inducing differentiation then death of melanoma cells, is unique among current melanoma therapies. Further evaluation is needed to assess other mechanisms that may be involved in tumor regression and determine whether anti-GDF6 mAb can be feasibly used in combination with other melanoma therapies, such as targeted BRAF inhibition or immunotherapy.

## MATERIALS & METHODS

### Cell lines and cell culture

A375, M14, HEK293T cells (ATCC) were maintained in DMEM supplemented with 10% FBS and 1% penicillin-streptomycin (Gibco). SK-MEL-28 cells were maintained in RPMI 1640 supplemented with 10% FBS and 1% penicillin-streptomycin (Gibco). All cell lines were kept at 37°C and 5% CO2. Cells cultured at the same time were pooled, counted, and then seeded in appropriate sized dish or plate. Dishes and plates were then subjected to designated treatments.

### Antibody production

Monoclonal antibodies recognizing GDF6 ligand were generated by injecting mice with a maltose binding protein-tagged GDF6 ligand (MBP-GDF6; amino acids 336-455 of the GDF6 proprotein). Hybridomas were produced, and antibodies from resulting hybridomas were screened by dot blot against MBP-GDF6.

### Western blots

Cells were lysed with ice-cold RIPA buffer containing a Complete protease inhibitor tablet (Roche). Protein concentration was measured using the Pierce BCA Protein Assay Kit (ThermoFisher). Samples were run on 10% polyacrylamide gels, transferred and developed using SuperSignal West Pico Plus (ThermoFisher).

Primary antibodies and dilutions used were: anti-GDF6 (Figure 1A,B Sigma PRS4691, 1:1000; otherwise this study, 1:500), anti-HA (Cell Signaling Technology 3724, 1:500), phospho-SMAD1/5/8 (Cell Signaling Technology 13820, 1:500), total SMAD1/5/8 (Cell Signaling Technology 6944, 1:500), alpha-tubulin (Cell Signaling Technology 3873, 1:1000). Secondary antibodies and dilutions used were: goat anti-rabbit IgG HRP-conjugated (Jackson ImmunoResearch 111-035-003, 1:2000), goat anti-mouse IgG HRP-conjugated (Jackson ImmunoResearch 115-035-003, 1:2000).

### Lentiviral infection and transfections

For expression of tagged GDF6, we PCR amplified GDF6 with primers including the sequence for hemeagglutinin (HA) and attB sites. We used gateway cloning to insert GDF6-HA into the pLenti CMV Hygro DEST (w117-1) vector. For stable gene knockdowns, a pLKO-1 lentiviral vector was used to deliver shRNA obtained from the RNAi Consortium through the UMMS RNAi core facility. One target sequence for GDF6 was used (GDF6.1, TRCN0000141818, target sequence: GCCAAGTGTTACATTGAGCTT). Virus was made using a second generation lentiviral packaging system in HEK293T cells and quantified using a p24 ELISA Kit (Takara Bio). Cells were infected with virus at a MOI of 2.5, with 8 ug/mL polybrene (Santa Cruz). Infection was performed as previously described^12^, except selection was made using 400 µg/mL of hygromycin for 10 days. Transfection of BMP luciferase was performed as previously described^12^. Briefly, pGL3 BRE Luciferase (Addgene) was prepared in a mixture with Lipofectamine 2000 (Invitrogen) in OptiMEM (Gibco) and added directly to A375 melanoma cells. Following 6 hours of transfection, pGL3 BRE Luciferase mixture was removed and replaced with DMEM growth media. Cells were allowed to recover for 18 hours and then prepared for assays as described below.

### In vitro drug treatments

Cells were cultured and plated as described above. Cells were allowed to adhere for 24 hours prior to treatment. For drug treatment with BMPi, stock solutions of DMH1 were prepared in DMSO and stored at - 20°C. Working solutions of 40 µmol of DMH1 were prepared in DPBS prior to treatment. DMSO was used for control treatment. For antibody treatment, antibodies were stored at 4°C and diluted to the appropriate concentration in PBS at the time of experiment. A non-targeting antibody was used for control treatment.

### Luminescence assays

Cells were prepared as described above and plated at a density of 1,500 cells/well in white-walled 96 well plates. Cells were treated with anti-GDF6 antibody or DMH1 24 hours after plating. Cell Titer Glo, Caspase 3/7 Glo, and Luciferase Assay System (Promega) were prepared per the manufacturers’ instructions. Following antibody and drug treatments for 24 hours, the appropriate assay reagent was added to each well, mixed, and read using 96-well plate luminometer (Perkin Elmer). Background luminescence was subtracted and replicate values were normalized to control treated wells.

### Mouse xenografts

Animals were handled in accordance with protocols approved by the UMMS IACUC. 6-8 week old athymic nude mice were subcutaneously injected with 5 X10^6^ SK-MEL-28 melanoma cells. Following engraftment, tumors were allowed to grow to a volume of 80-100 mm^3^. Tumor volumes were made from external caliper measurements and calculated using the formula V = (W x W x L)/2, where W is the shortest width and L is the longest length of the tumor. After reaching 80-100 mm^3^, mice were randomized to experimental and control groups then treated with designated compound. For treatment with anti-GDF6 antibody, two groups of 6 mice each received biweekly doses of 10 mg/kg of anti-GDF6 antibody or control non-targeting IgG antibody (Bioxcell BP0087) via intraperitoneal injection.

### Immunohistochemistry

From mouse xenografts with human melanoma cells, formalin-fixed, paraffin-embedded tissues were processed to obtain 5µm sections. Sections were stained with H&E, cleaved Caspase-3 (Cell Signaling Technology 9664; 1:100), Ki-67 (Dako M7240, 1:100) and evaluated, anti-GDF6 (this study, 1:250). The numbers of cleaved Caspase-3 or Ki-67-positive cells were counted manually, and the total number of cells in each field was calculated using ImageJ software. Sections were also stained with phospho-SMAD (Cell Signaling Technology 13820, 1:50) and evaluated. Phospho-SMAD-stained sections were scored for immunointensity (0-4) and immunopositivity (0-3), which were then multiplied.

### Flow cytometry

A375 melanoma cells incubated with HA-tagged GDF6 or A375 melanoma cells expressing HA-tagged GDF6 were treated with Alexa Fluor 488-conjugated anti-HA antibody. Flow cytometry was performed using a BD Aria II instrument. Flow cytometry data were analyzed using FloJo software.

### Immunofluorescence

A375 cells were grown directly on collagen IV-coated coverslips (Sigma), fixed in 3.7% formalin, permeabilized using 0.1% Triton X-100, and treated with 0.1% SDS. They were blocked in 1% BSA and then incubated with primary antibody diluted in blocking solution in a humidity chamber at 4°C overnight, washed with 1x PBS, and incubated with secondary antibody in blocking solution. Cells were mounted using mounting media containing DAPI (Vector Laboratories). All secondary antibodies were Alexa Fluor conjugates (488 and 647) (Thermofisher) and used at a 1:500 dilution. Primary antibodies and dilutions used were: anti-GDF6, 1:500 (this study), phospho-SMAD1/5/8, 1:500 (Cell Signaling Technology 13820), cleaved Caspase-3 (Cell Signaling Technology 9664; 1:250), Ki-67 (Dako M7240, 1:250).

### RT-qPCR

RNA was isolated from A375 cells using the RNeasy kit (Qiagen) per manufacturer’s protocol. cDNA was synthesized from purified RNA using the SuperScript III First Strand Synthesis kit (ThermoFisher). Reaction mixes were assembled with SYBR Green RT-PCR master mix (ThermoFisher), primers, and 25ng cDNA, and analyzed using a QuantStudio 3 Real-Time PCR System. All samples were normalized to *β-Actin*, and fold changes were calculated using the ΔΔCt method.

## SUPPLEMENTARY FIGURE LEGENDS

**Supplementary Figure 1. Identification of anti-GDF6 antibodies.** (A) Dot blots to test reactivity of hybridomas. MBP-GDF6 was blotted as the target protein. (B) Western blots of selected hybridomas on GDF6 protein isolated from A375 melanoma cell lysates showing recognition of full-length 55 kDa pro-GDF6 and 15 kDa mature GDF6. (C) Assessment of viability using CellTiter Glo of A375 melanoma cells following 24 hours of treatment with purified anti-GDF6 mAbs. n = 3 technical replicates.

**Supplementary Figure 2. Anti-GDF6 mAb reduces phospho-SMAD1/5/8 protein.** (A) Western blot of protein from A375 cells treated with anti-GDF6 antibody and control IgG antibody. (B) Quantification of Phospho-SMAD1/5/8 to total SMAD1/5/8 ratios from western blots.

**Supplementary Figure 3. Tumor volumes in mice treated with anti-GDF6 antibody.** Tumor volume growth curves of individual mice treated with anti-GDF6 mAb (left) and control IgG mAb (right). Treatments are indicated by arrows. P indicates a tumor that was below the measureable limit but palpable. Open circles indicate mice sacrificed for on-treatment tumor analyses. X mark indicates a mouse that died after the treatment window but prior to the end of study.

