## Supplementary figures and images for "INHIBITION OF LIGAND-DEPENDENT BMP SIGNALING BLUNTS MELANOMA GROWTH"

### Supplemental Figure 1

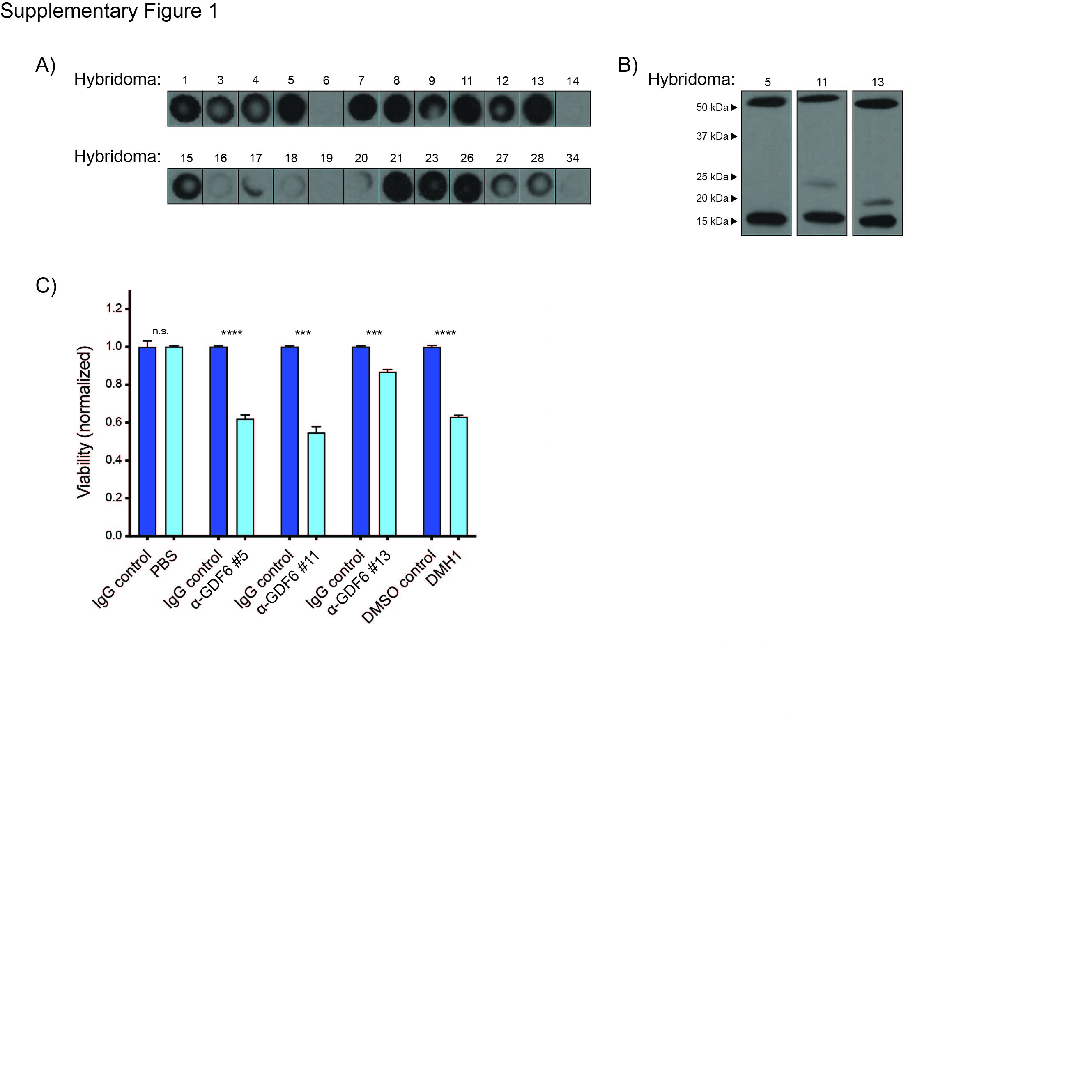

### Supplemental Figure 2

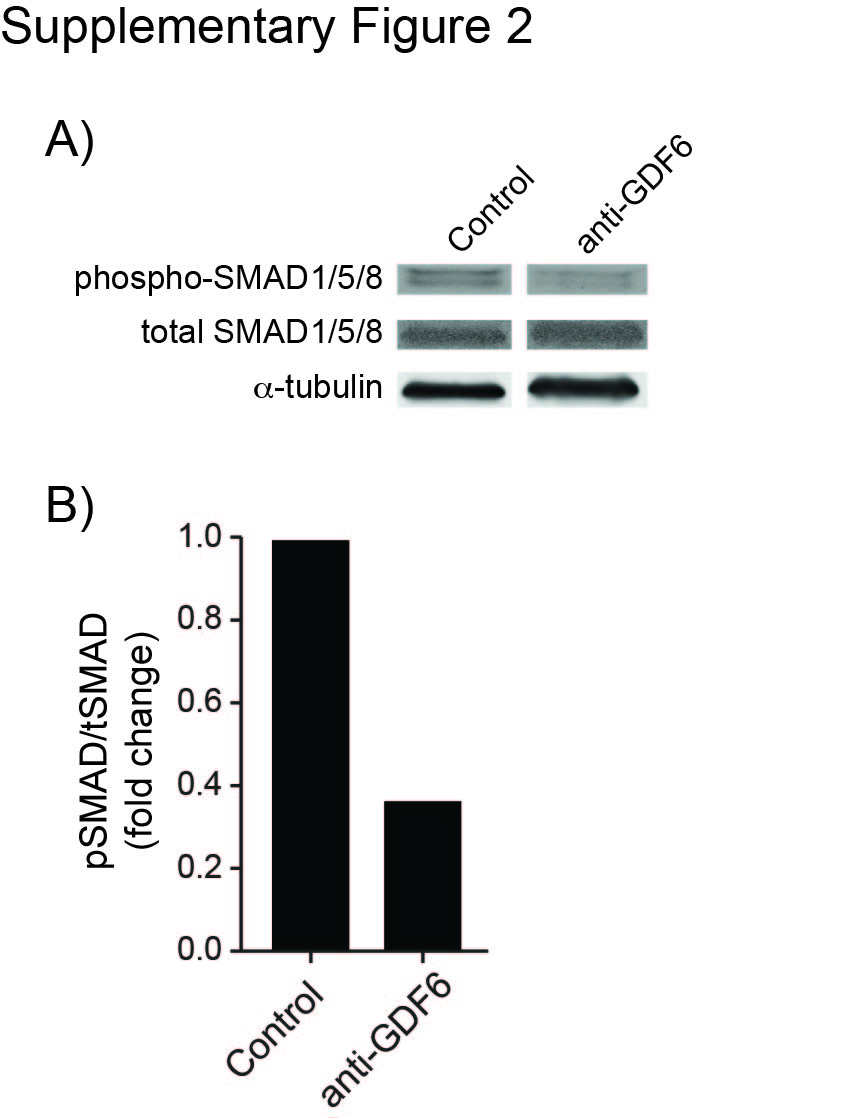

### Supplemental Figure 3

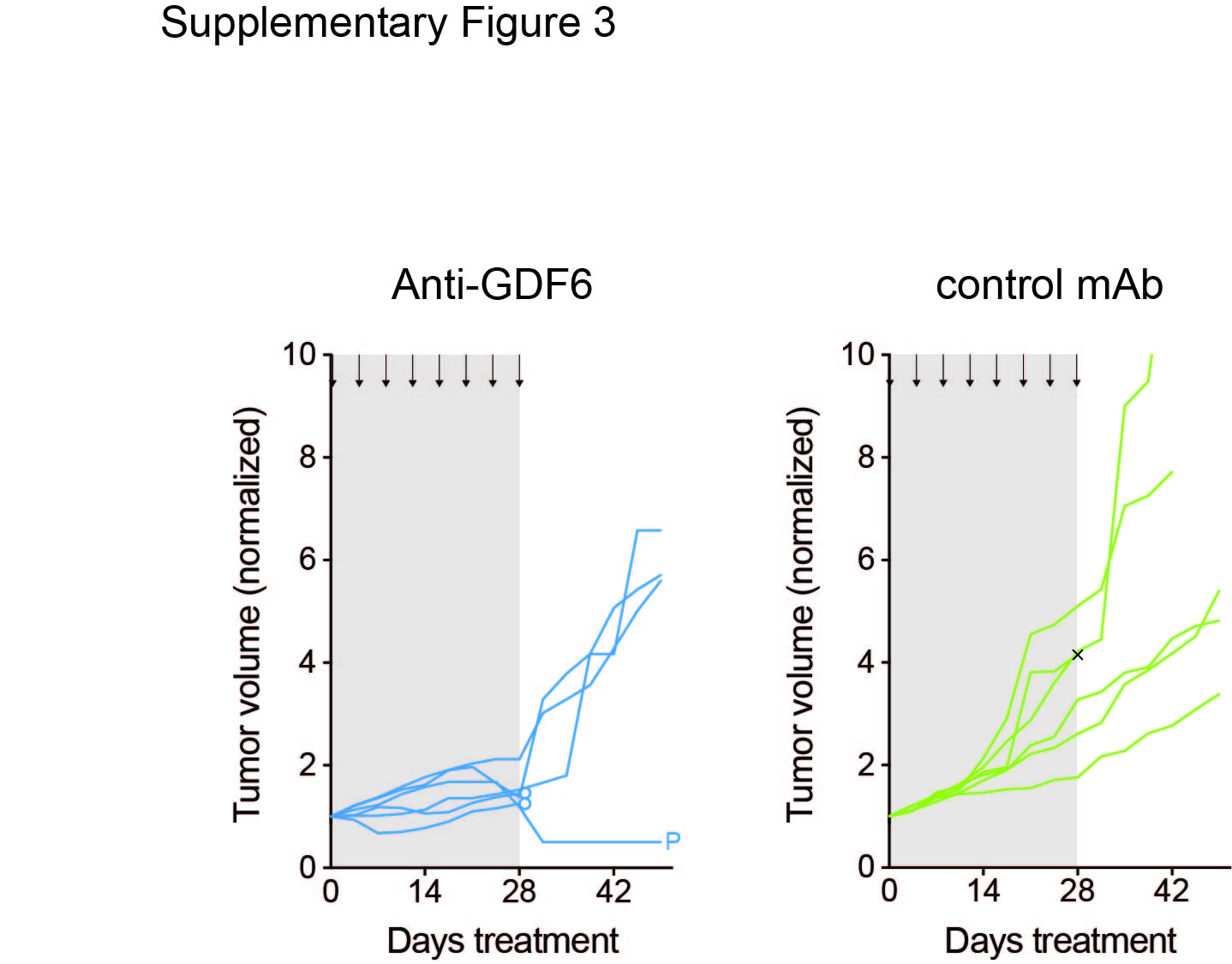
